# Accurate differential analysis of transcription factor activity from gene expression

**DOI:** 10.1101/296210

**Authors:** Viren Amin, Murat Can Cobanoglu

## Abstract

We present EPEE (Effector and Perturbation Estimation Engine), a method for differential analysis of transcription factor (TF) activity from gene expression data. EPEE addresses two principal challenges in the field, namely incorporating context-specific TF-gene regulatory networks, and accounting for the fact that TF activity inference is intrinsically coupled for all TFs that share targets. Our validations in well-studied immune and cancer contexts show that addressing the overlap challenge and using state-of-the-art regulatory networks enable EPEE to consistently produce accurate results. (Accessible at: https://github.com/Cobanoglu-Lab/EPEE)

## Main text

Differential analysis of gene expression data is commonly used to dissect the mechanism of phenotypes of interest^1^. Commonly used differential expression (DE) methods^2–5^ do not account for the regulation of gene expression by transcription factors (TFs), even though TFs have a key role in controlling the transcriptome^6^. This shortcoming has prompted the development of differential regulation (DR) methods^7–13^, however these methods either do not permit the full representation of available context-specific regulatory data^7,8,10–13^ or interrogate regulators individually^9,14^ resulting in potential false positives for overlapping regulons^8^. To address these issues, we have developed the Effector and Perturbation Estimation Engine (EPEE) which infers the regulatory activity of all TFs jointly, constrained under the context-specific TF regulatory networks (**Fig. 1**).

**Figure 1.**
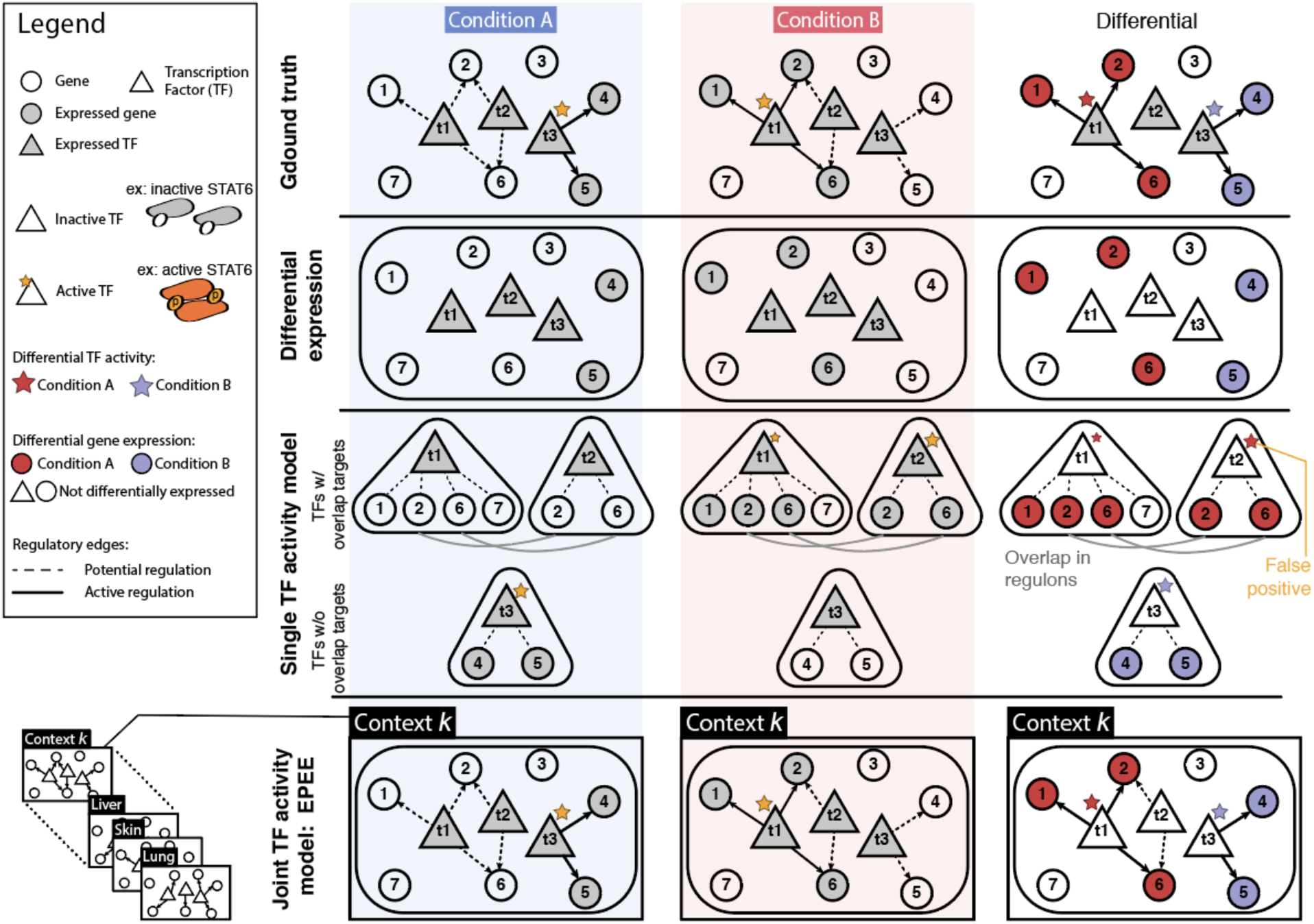
Schematic overview of EPEE in relation to alternative approaches. We construct a schematic example to illustrate our motivation for developing EPEE. In this schematic, the ground truth (top row) is that one TF is active in each condition. Differential expression methods (second row) do not account for any regulatory relationship among genes, and simply report the genes with different expression. Single TF activity model based differential regulation methods (third row) evaluate each TF individually, and this is problematic when TF regulons overlap, leading to false positives. EPEE (bottom row) uses the appropriate context-specific TF-gene regulatory network and models all TF activity as a single multivariate regression problem, to address both context-specific changes and overlapping regulons.

EPEE models gene expression as the result of latent activity by TF gene products. We use the context-specific TF regulatory graph to prune inaccessible targets for each TF regulon, and then infer the activity of all TFs jointly with a single multivariate model (**Supplementary Note 1**). The activity of any TF over its target genes are inherently related with each other. We use graph constrained fused lasso^15^ to reflect that intuition. Briefly, fused lasso^16^ is a method for multivariate regression where the inference of model parameters are “fused” according to *a priori* established relationships. Graph constrained fused lasso^15^ is an extension to the setting where the inferred parameters are related with a graph structure. In EPEE, we fuse the parameters of the model to respect the context-specific regulatory network during TF activity inference. Crucially, since we represent the inference of all TF activity as a single (multivariate regression) problem, our inference automatically adjusts for overlapping TF regulons.

Overlap among TF regulons is widely prevalent, based on the state-of-the-art TF regulatory networks^17^ (**Supplementary Fig. 1**). Of the 206,403 TF-TF pairs in the CD4^+^ T cell context, 206,352 (99.9%) overlap with each other to some degree (**Supplementary Fig. 1c**). To check if this is a feature specific to a few contexts, we evaluated all the 394 human contexts for which TF regulatory graphs are available^17^. We found that even the least complex context has overlaps among 96.4% of all the TF-TF pairs, whereas the median was 98% (**Supplementary Fig. 1d**). Causal inference theory dictates that all latent random variables that cause an effect become dependent when conditioned on the outcome^18^. Consequently, when inferring TF activity (latent cause) from gene expression (observed outcome), the activity inference for all TFs with overlapping regulons are dependent on each other. Hence, we argue that the latent TF activity inference must be solved jointly and we propose a single, multivariate model.

Most previous approaches on TF activity inference calculate each TF’s activity individually^7–14^ and only aggregate at a later stage. For example, in the popular Gene Set Enrichment Analysis^14^ (GSEA), the running sum statistic evaluates each TF’s activity individually. Likewise, MARINa^8^ or VIPER^12^ use mutual information to estimate TF activity, and this measure evaluates each TF separately. These two latter methods^8,12^ utilize a *post hoc* correction for this problem, called “shadow analysis”, but its generality is unclear. To demonstrate the improvement that the joint TF activity model in EPEE provides over previous approaches, we conducted comparative evaluations.

We conducted comparative validation studies in two independent and well-studied contexts with known driver TFs: T helper cell differentiation and colorectal adenocarcinoma. The immune cell context enables us to compare methods within the context of normal transcriptional regulation. The cancer dataset, on the other hand, enables us to evaluate performance under the disrupted regulatory state that accompanies genomic instability in carcinogenesis^19^. Furthermore, the T cell data is homogeneous (i.e. highly purified biological replicates of the exact same cell types), while the cancer data from TCGA^20^ represents heterogeneous input because of both tumor purity^21^ and subclonality^22^. The final major difference between the two datasets is the number of samples: the immune cell differentiation dataset has five samples per class (representative for a standard research laboratory effort), whereas the cancer dataset has close to five hundred samples (representative of a major consortium effort). In summary, we carefully selected these two validation studies to cover the diverse range of conditions in which differential TF activity assessments can be used.

Our first validation study was to identify the known driver TFs controlling CD4^+^ naïve T cells differentiation to T helper cells 1, 2, and 17 (T_h_1, T_h_2, T_h_17). We selected the driver TFs as follows: STAT6^23–27^ and GATA3^28^ for T_h_2; TBX21^29^, STAT1^30^, STAT4^30^ for T_h_1; and RORα^31^, STAT3^31,32^, and ARID5A^33^ for T_h_17 differentiation of CD4^+^ T cells. We input the same RNA-seq data^34^ to all methods (**Supplementary Note 2**). We quantified performance as the ranking of the ground truth TFs with each method, with higher rank signifying better performance. We used ten alternative methods that represent a variety of inference and regulation models (**Supplementary Note 3**). Eight were developed and/or previously used to estimate regulatory activity^7–14^. We also included two differential expression methods to test the effectiveness for searching for an increase in the expression of the TF itself as a marker of its activity: ANOVA as a canonical differential expression analysis method^11^, and sleuth as a state-of-the-art DE method^5^. Of the eight differential regulation methods, three do not require an explicit regulatory network^7,10,11^ while the remaining five utilize a mathematical representation that evaluates each TF regulon individually. Among the alternatives, DISCERN^11^ is the only method that also solves the TF activity inference problem jointly for all TFs, albeit cannot incorporate a fully specified TF-gene network. Therefore, we tested multiple members of each category of method a practitioner can use to identify TF activity. We observed that while other methods show significant variation and often fail to properly identify the known drivers, EPEE is consistently accurate in ranking ground truth TFs as differentially active **(Fig. 2)**.

**Figure 2:**
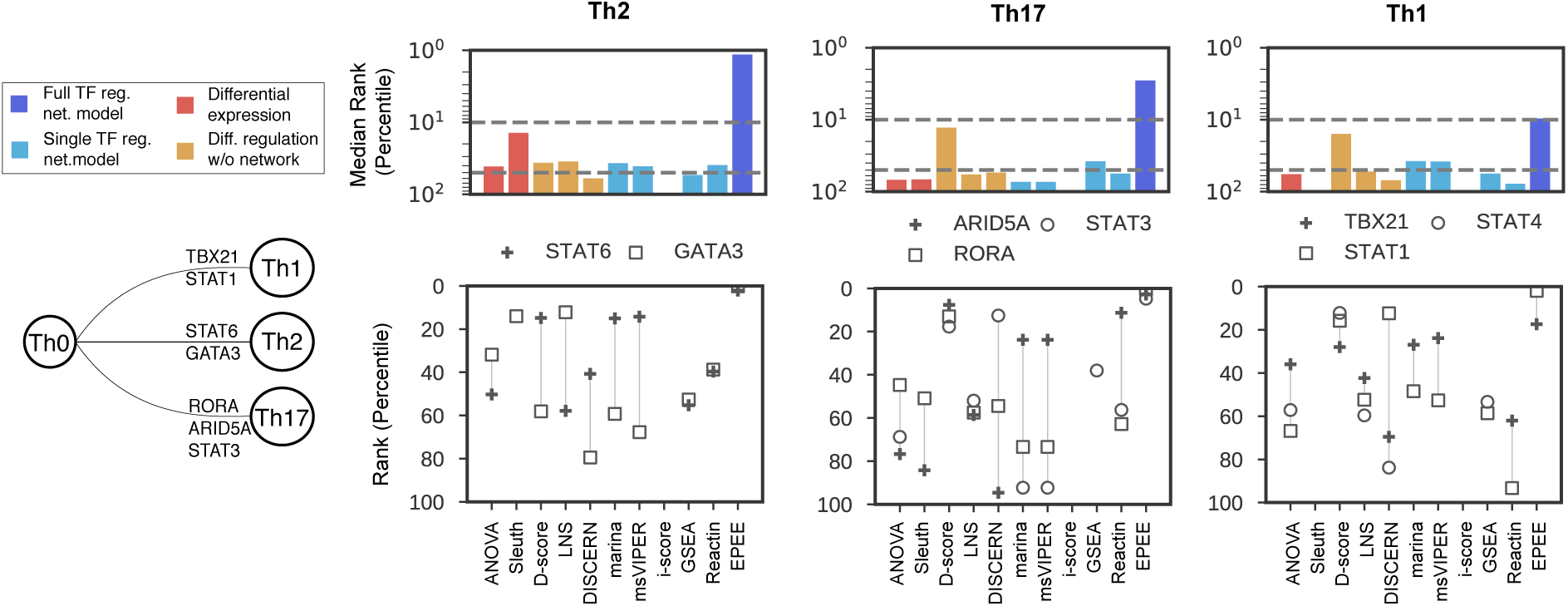
EPEE infers transcriptional regulators reliably and effectively. We tested regulator inference in CD4 naïve T cell differentiation to T helper 1 (T_h_1), T_h_2, T_h_17 cells. Drivers are known for each pathway therefore we can compare performance. EPEE performs remarkably better than all other DE (red) and DR (gold: no network, light blue: transcriptomic data driven network, dark blue: full regulatory network) methods. In the log percentile plots, the top dashed line represents the 90th percentile threshold, the bottom dashed line represents the 50the percentile.

EPEE can incorporate context-specific TF regulatory networks with weighted edges, which are available thanks to recent large-scale data collection efforts such as ENCODE^35^ or FANTOM5^36^ (**Supplementary Table 2**). Among the alternative methods, only REACTIN^9^ and i-score^13^ can utilize these rich regulatory graphs. GSEA uses MSigDB^37^ TF target gene sets, that cannot represent TF-gene edge weights; MARINa and VIPER utilize ARACNE^38^ networks built using only transcriptomic data; while D-score^7^, LNS^10^, and DISCERN^11^ do not utilize any regulatory graph. The methods without networks serve as controls for the quality of the state-of-the-art regulatory networks: if the current networks are too deficient (not all TF motifs are available, for example) then the network-independent methods should perform better, and *vice versa*. For the methods that can utilize context-specific TF regulatory graphs, including our method, we input the CD4^+^ T cell network that Marbach and colleagues curated^17^ using FANTOM5^36^ data. EPEE consistently outperformed methods with a range of network inputs, showing that EPEE provides an improved model to benefit from the state-of-the-art regulatory networks.

We then elucidated the contributions to our method’s performance. The regularization provides consistency (**Supplementary Fig. 2**), while the context-dependent network contributes to the accuracy (**Supplementary Fig. 3**). EPEE can resolve TFs with high regulon overlap, while other methods cannot necessarily do so (**Supplementary Fig. 4**). Furthermore, the performance is not the result of parameter tuning. We used an entirely independent dataset of acute myeloid leukemia (AML) gene expression (microarray data) to determine the default hyperparameter values (**Supplementary Fig. 5**) and we simply used these default settings for all the results in our study. Likewise, for the competing methods we also used the default settings, unless suggested otherwise in the manual or documentation for that method (**Supplementary Note 3**).

**Figure 3:**
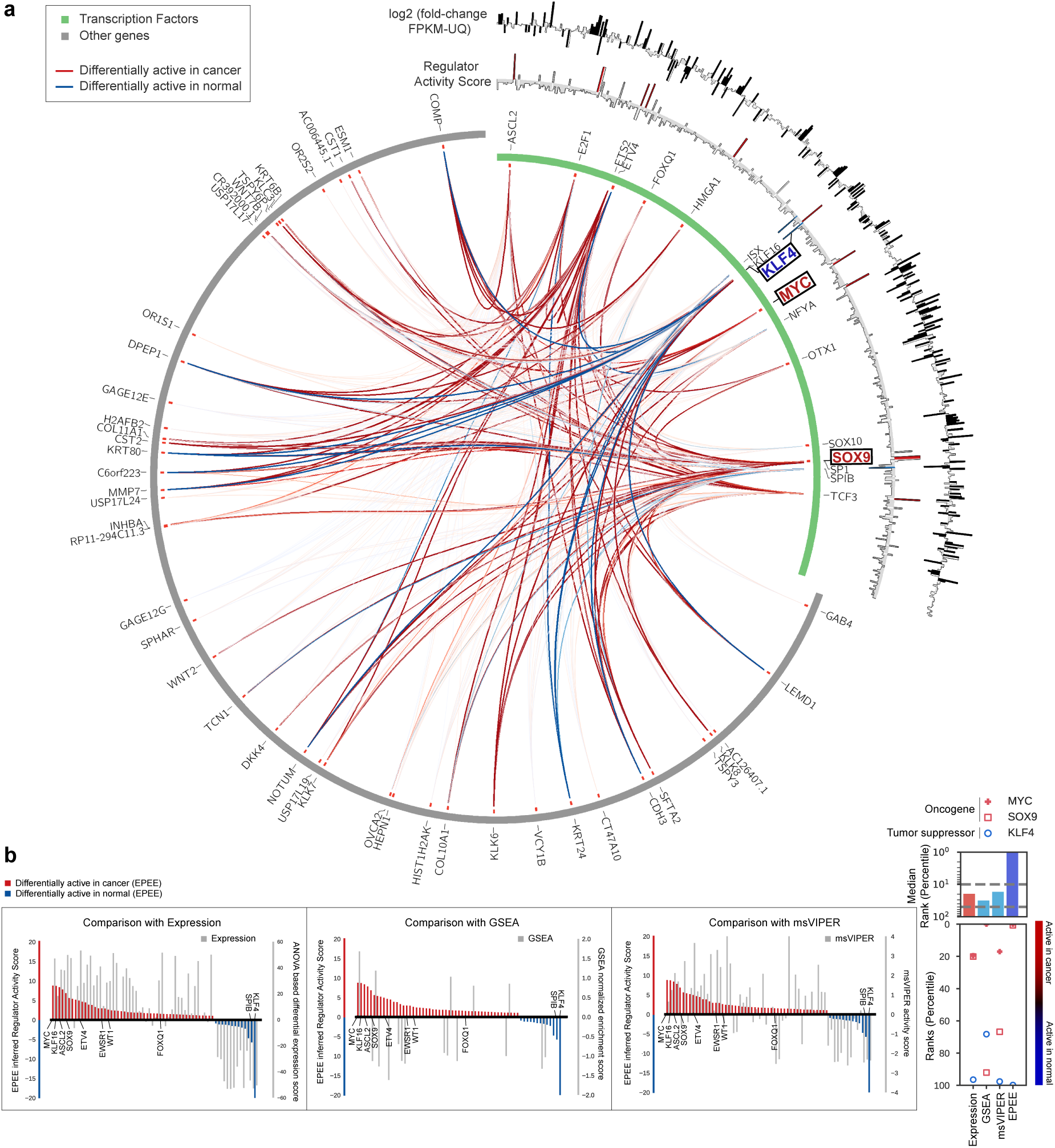
EPEE can accurately infer differential regulation events in cancer. (a) We conducted differential analysis of TF activity between colorectal adenocarcinoma (COAD) samples and tissue-matched controls using TCGA data. Circos plot shows differential regulation with red and blue TF-gene edges having high activity in cancer and normal. Green band maps TFs and grey band maps perturbed genes. (b) We also performed the same analysis using GSEA, msVIPER, and ANOVA. EPEE accurately identified the oncogenes MYC and SOX9 as differentially active, ranking MYC as the most differentially active TF in cancer. EPEE also identified tumor suppressor KLF4 to be differentially active in normal.

To test whether EPEE can generalize to other biological domains, we applied EPEE to identify driver TFs from colorectal adenocarcinoma (COAD) gene expression data in TCGA (**Fig. 3)**. In this context, we compared against msVIPER and GSEA due to their popularity, and ANOVA as the standard differential expression method. We used known oncogenes MYC^20,39^ and SOX9^20,40^ as ground truth TFs that are differentially active in cancer. On the other end, we used KLF4^41–43^ as a TF known to be differentially active in normal tissue, since KLF4 is commonly inactivated in cancer via diverse mechanisms such as miRNA silencing^44,45^. We used EPEE to identify both the statistically significantly perturbed genes and regulators (Benjamini-Hochberg FDR < 0.05, based on permutation tests as described in **Supplementary Note 1**) and show the inferred TF-gene regulation changes (**Fig. 3a**). EPEE correctly identified the differential activity of all three TFs (**Fig. 3b**). GSEA failed to identify SOX9 as an oncogene, and KLF4 as differentially active in normal. msVIPER correctly identified KLF4 as differentially active in normal, but misplaced both MYC and SOX9 by inferring many other TFs to be more active in cancer. Overall, EPEE outperformed alternative methods in cancer as well as it did in immune cells.

Finally, we used EPEE to infer differential TF activity separately for each consensus molecular subtype (CMS) of colorectal adenocarcinoma (**Supplementary Fig. 6**). We found that MYC, SOX9 and KLF4 activity were homogenous across each CMS subtype despite varying purity estimates in each CMS (**Supplementary Fig. 7**). We also discovered subtype-specific TFs with differential activity, some of which were reported to be colorectal adenocarcinoma related (ASCL2^46–49^, ETV4^50^, HSF1^51^) while others represent novel predictions that can be tested by follow-up experimental work.

In conclusion, we assert that the prevalent overlap among TF regulons causes problematic TF activity inference by readily available existing methods and thus present a novel solution that models all TF activity as a single multivariate regression problem. We demonstrate consistently accurate results on well-studied immune and cancer contexts. We provide our method EPEE open source and freely available.

## Acknowledgements

We would like to acknowledge Curtis Thorne, Didem Ağaç, David Farrar, Anne Satterthwaite, Maxim Grechkin, and Daniel Marbach for their helpful discussions.

